# Adapting a two-photon scanning microscope for simultaneous single-photon imaging of an infrared dopamine sensor

**DOI:** 10.64898/2026.01.13.699388

**Authors:** Matthew Tarchick, Franklin Caval-Holme, Ben Smith, Petra Mocellin, Markita Landry, Natsumi Komatsu, Marla B. Feller

## Abstract

We describe a novel method for adapting a two-photon scanning microscope to enable simultaneous detection of two-photon generated visible fluorescence and single-photon generated near-infrared (nIR) fluorescence. In this configuration, nIR fluorescence is routed through a single-mode optical fiber before detection by a photomultiplier tube. This fiber coupling offers two advantages: first, the optical fiber functions as a pinhole aperture, allowing for improved optical sectioning of the nIR signal; second, it minimizes nIR background fluorescence. To validate the effectiveness of this design, we conducted two sets of experiments. First, we compare two fluorescence indicators of the neurotransmitter dopamine: the genetically encoded indicator GRAB_DA_ and single walled carbon nanotube based optical nanosensors (nIRCats). Although nIRCats exhibit lower affinity for dopamine than GRAB_DA_, this property allows for identification of high concentration release sites in the striatum. Second, we simultaneously imaged depolarization-induced calcium changes and dopamine release in the retina. Together, these results demonstrate the utility of integrating confocal nIR detection into a two-photon platform for simultaneous functional imaging across complementary spectral channels.

## Introduction

The neuromodulator dopamine plays a critical role in multiple brain functions (Speranza et al. 2021). Understanding the mechanisms by which dopamine impacts brain activity requires measuring spatial and temporal dynamics of extracellular dopamine concentrations (Sippy and Tritsch 2023). For example, diverse firing patterns in the ventral midbrain modulate distinct brain functions: tonic firing in substantia nigra dopaminergic neurons produces sustained elevation of dopamine in the dorsal striatum that promote a permissive environment for movement (Da Silva et al. 2018), while burst firing in the ventral tegmental area generates transient elevations of dopamine associated with reward prediction errors (Bayer and Glimcher 2005). In the retina, dopamine similarly acts as a critical neuromodulator, regulating short-term light adaptation, modulating receptive field properties, and contributing to circadian changes in retinal function (Witkovsky 2004; Roy and Field 2019; Jackson et al. 2012).

To characterize the dynamics of dopamine release, the field has turned to optical sensors. One popular class of these sensors include genetically encoded sensors in which green fluorescent proteins (GFP) is integrated with dopamine receptors - (dLight and GPCR-activation-based dopamine sensors GRAB_DA_) (Sun et al. 2018; Patriarchi et al. 2018). These sensors have proven to be powerful detectors of dopamine release but require viral transduction of the sensors and therefore are limited to use in genetically tractable organisms.

Another tool for detecting dopamine optically is synthetic near-infrared (nIR) catecholamine nanosensors (nIRCat) (Beyene et al. 2019; Del Bonis-O’Donnell et al. 2021). nIRCat nanosensors are synthesized from single-walled carbon nanotubes, which fluoresce in the near-infrared, a window that prevents scattering in tissue. They are functionalized with (GT)₆ DNA sequences to have specificity for dopamine and norepinephrine over others. nIRCat sensors have a lower affinity for dopamine than the genetically encoded sensors (Beyene et al. 2019), but they allow for the detection of hot spots of dopamine release, such as in slices of the dorsal striatum. In addition, nIRCats have been incorporated into films, allowing for the characterization of dopamine release along isolated dopaminergic axons (Bulumulla et al. 2022; Elizarova et al. 2022).

Furthermore, nIRCat’s near infrared fluorescence is spectrally well-separated from visible fluorophores, providing an opportunity to study dopamine release simultaneously with calcium dynamics or other neurotransmitter activities with visible fluorophores. Dual imaging of nIRCat and visible wavelength fluorophores has been demonstrated (Bulumulla et al. 2022; Elizarova et al. 2022); however, visible wavelength fluorophores were used to label neurons in those studies, rather than to monitor real-time activity.

Here we introduce a dual-modal scanning microscope that integrates two-photon excitation of a visible fluorophore with single photon excitation of nIR nanosensors. Optical channels are spectrally resolved and independently coupled to maximize signal fidelity. We validated our design by measuring real-time dopamine dynamics in the striatum with simultaneous imaging of nIRCats and GRAB_DA3m_. We further demonstrate real-time imaging of depolarization-induced calcium transients and simultaneous dopamine release in the retina. While we focus on monitoring sensors of Ca^2+^ and neurotransmitter release as a proof of concept in this study, this framework generalizes to any visible fluorophores, and supports flexible expansion to other analytes or modalities.

## Materials and Methods

### Experimental model

All experiments were performed in adult male and female C57Bl6 mice (Jackson Laboratory). The animals were maintained on a 12:12 light cycle (lights on at 07:00). All animal procedures were performed in accordance with the University of California Berkeley Institutional Animal Care committee’s regulations and Use Committees and conformed to the National Institutes of Health’s Guide for the care and use of laboratory animals, the Public Health Service Policy, and the Society for Neuroscience Policy on the Use of Animals in Neuroscience Research.

### Stereotactic viral injection

The injections were performed under general ketamine– dexmedetomidine anesthesia using a stereotaxic instrument (Kopf Instruments, Model 1900) as previously described (Lammel et al. 2008). For the GRAB_DA_ experiment, 400 nl of AAV2/9-hSyn-GRAB_DA3m-WPRE-hGH polyA (Brain VTA, PT-4720) were injected bilaterally in the dorsal striatum. The coordinates used to target the dorsal striatum were AP: +1; ML: +/- 1.5; DV: -3.0 relative to bregma and based on the Allen Mouse Brain Atlas. After all injections the needle was left in place for at least 5 minutes before withdrawal. The animals were kept on a heating pad until recovered from anesthesia. Experiments were performed 3-4 weeks after stereotactic injection.

### Acute brain slice preparation

3-4 weeks after viral injection mice were deeply anaesthetized with pentobarbital (200 mg/kg ip; Vortech). After intracardial perfusion with ice-cold artificial cerebrospinal fluid (aCSF containing 119 mM NaCl, 26.2 mM NaHCO_3_, 2.5 mM KCl, 1 mM NaH_2_PO_4_, 3.5 mM MgCl_2_, 10 mM glucose, and 0 mM CaCl_2_) the brain was rapidly extracted and sliced with a Leica VT1200 S vibratome. Before use, slices were incubated at 37°C for 60 min in oxygen-saturated aCSF (119 mM NaCl, 26.2 mM NaHCO_3_, 2.5 mM KCl, 1 mM NaH_2_PO_4_, 1.3 mM MgCl_2_, 10 mM glucose, and 2 mM CaCl_2_) and then transferred to room temperature for 30 min before nIRCat labeling and imaging experiments.

### Retina preparation

Mice were deeply anesthetized with isoflurane inhalation and killed by decapitation. Eyes were immediately enucleated, and retinas were dissected in oxygenated (95% O_2_, 5% CO_2_) aCSF (in mM as follows: 119 NaCl, 2.5 KCl, 1.3 MgCl_2_, 1 K_2_HPO_4_, 26.2 NaHCO_3_, 11 D-glucose, and 2.5 CaCl_2_) at room temperature under white light. In some experiments, 0.1 mM sulforhodamine 101 (SR101, Invitrogen) was added for visualization of vasculature. Each isolated retina was cut into two pieces. Each piece of retina was mounted over a 1-2 mm^2^ hole in nitrocellulose filter paper (Millipore) with the photoreceptor layer side down, dark-adapted for 1 h, and transferred to the recording chamber of a two-photon microscope for imaging. The whole-mount retinas were continuously perfused (3 ml/min) with oxygenated aCSF warmed to 32°-34°C by a regulated inline heater (TC-344B, Warner Instruments) for the duration of the experiment. Additional retina pieces were kept in the dark at room temperature in aCSF bubbled with 95% O_2_, 5% CO_2_ until use (maximum 8 h).

### nIRCat synthesis

HiPCo SWCNT slurry (NanoIntegris) and ssDNA (Integrated DNA Technologies) were mixed with a final NaCl concentration of 10 mM. The mass ratio of DNA to SWCNT was 2:1. The DNA sequence was GTGTGTGTGTGT. The mixture was probe-tip sonicated (Cole-Parmer Ultrasonic Processor, 3-mm tip) in ice-cold water for 30 min at 50% amplitude. The resulting suspensions were centrifuged at 21,000 g for 4 hours at 4°C and the supernatant was collected. Nanosensors were diluted to 200 mg/L in 10 mM NaCl and stored at 4°C.

### nIRCat labeling (brain slices)

aCSF (119 mM NaCl, 26.2 mM NaHCO_3_, 2.5 mM KCl, 1mM NaH_2_PO_4_, 1.3 mM MgCl_2_, 10 mM glucose, 2 mM CaCl_2_, all purchased from Sigma-Aldrich) was prepared and bubbled with carbogen gas (oxygen/carbon dioxide 95% O_2_, 5% CO_2_, Praxair). To label the slice with nIRCat, brain slices containing striatum were transferred to an incubation chamber filled with carbogen-bubbled aCSF at room temperature. nIRCats were applied to the surface of brain slices with a pipette to a final concentration of 2 mg/L and incubated for 15 min. Slices were rinsed for 5 sec with bubbled aCSF through 3 wells of a 24-well plate to remove unlocalized nanosensors, and rested for at least 15 min before imaging.

### nIRCat and Cal 520 labeling (retina)

Retinas were transferred to the recording chamber of a two-photon microscope. nIRCat sensors (10 mg/mL in ACSF) were combined 1:1 with Cal 520 AM (AAT Bioquest) and bulk-loaded into the retina using a multicell bolus loading technique (Blankenship et al. 2009; Stosiek et al. 2003). Bolus loading was targeted to the retina’s inner-plexiform layer so as to deposit the sensor near processes of dopaminergic amacrine cells. Retinas were allowed to equilibrate for one hour before imaging.

### Imaging

Carbogen-bubbled aCSF was flown through the microscope chamber. Slices labeled with nIRCats were placed in the chamber with a tissue harp, and a glass pipet or a bipolar stimulation electrode (MicroProbes for Life Science Stereotrodes Platinum/Iridium Standard Tip) was positioned to the targeted field of view, which was adjusted with a 4x objective followed by a 60x objective (Olympus 1.00 NA LUMPlanFLN). The microscope was controlled by ScanImage Software. Fluorescent images were acquired at frame rates of 1.47 frames/s in a 130 µm^2^ (256x256 pixel) field of view. An ultrafast pulsed laser (tuned to 950 nm) provided fluorescence excitation.

To measure depolarization-evoked dopamine release, 1 second of 100 mM potassium solution or 1 millisecond of 0.1 to 0.3 mA stimulation was applied, and this stimulation was repeated at least three times with the same field of view with 5 minutes between each stimulation. For the pharmacological measurements, 10 µM of quinpirole (Fisher Scientific, 10-611-0) was applied to the chamber through aCSF perfusion and slices were incubated for 15 minutes before imaging. Images were acquired in the same field of view as the one before quinpirole application.

### Image analysis for an infrared dopamine sensor

Imaging movie files were processed using a Python script (https://github.com/NicholasOuassil/NanoImgPro). Fluorescent modulation ΔF/F_0_ was calculated as ΔF/F = (F-F_0_) /F_0_, where F_0_ is the average intensity before the stimulation and F is the dynamic fluorescence intensity from the entire field of view. Next, a 8x8 pixel (corresponding to 6 µm by 6 µm) grid mask was applied to the image stack to minimize bias and improve stack processing time. For each grid square ΔF/F_0_ was calculated, and they were identified as regions of interest (ROI) if the ΔF/F_0_ around time of stimulation is 2 standard deviations above the baseline activity. All the parameters reported in this study are averaged over at least three stimulations.

### Image analysis - retinal Ca^2+^ and nIRCat imaging

For each field of view, a single ROI was defined and fluorescence data extracted from both spectral channels in FIJI. ROIs encompassed the entire field of view, or the region labeled with nIRCat sensor. Wave-triggered and K^+^ puff-triggered average responses were computed using a Python script (https://github.com/FellerLabCodeShare/nIR-catecholamine-sensor). Fluorescence signals were low-pass filtered at 0.68 Hz using a third-order Butterworth filter. Fluorescent modulation ΔF/F_0_ was calculated as ΔF/F = (F-F_0_) /F_0_, where F_0_ is the median intensity during the imaging trial. In some cases, nIRCat fluorescence exhibited an exponential decay during the first 30-60 sec. of imaging. These decays were removed by fitting and then subtracting from the signal a double exponential. Spontaneous retinal waves were identified by Z-scoring the Cal 520 signal and finding peaks that exceeded the mean by >3 standard deviations and were separated by at least 10 sec. Wave and puff-triggered averages were computed within a 60 sec. window centered on the frame prior to the wave peak or K^+^ puff. Responses to puffs and waves were measured from the triggered averages by computing the difference between the peak values (a mean within a 3 sec. window centered on the peak) in 10 sec. windows before and after the event trigger.

## Results

### Integrating an infrared detection pathway into a scanning two-photon microscope

nIRCat sensors fluoresce in the near-infrared (Figure 1A). Thus far, imaging of these sensors in brain tissue have been based on epifluorescent imaging, using InGaAs cameras that are sensitive to long wavelengths. Here we present a strategy for integrating an infrared detection pathway into a two-photon (2P) laser scanning microscope (tuned 920-950 nm), which will allow for multiplexed imaging of these sensors with visible wavelength sensors.

**Figure 1:**
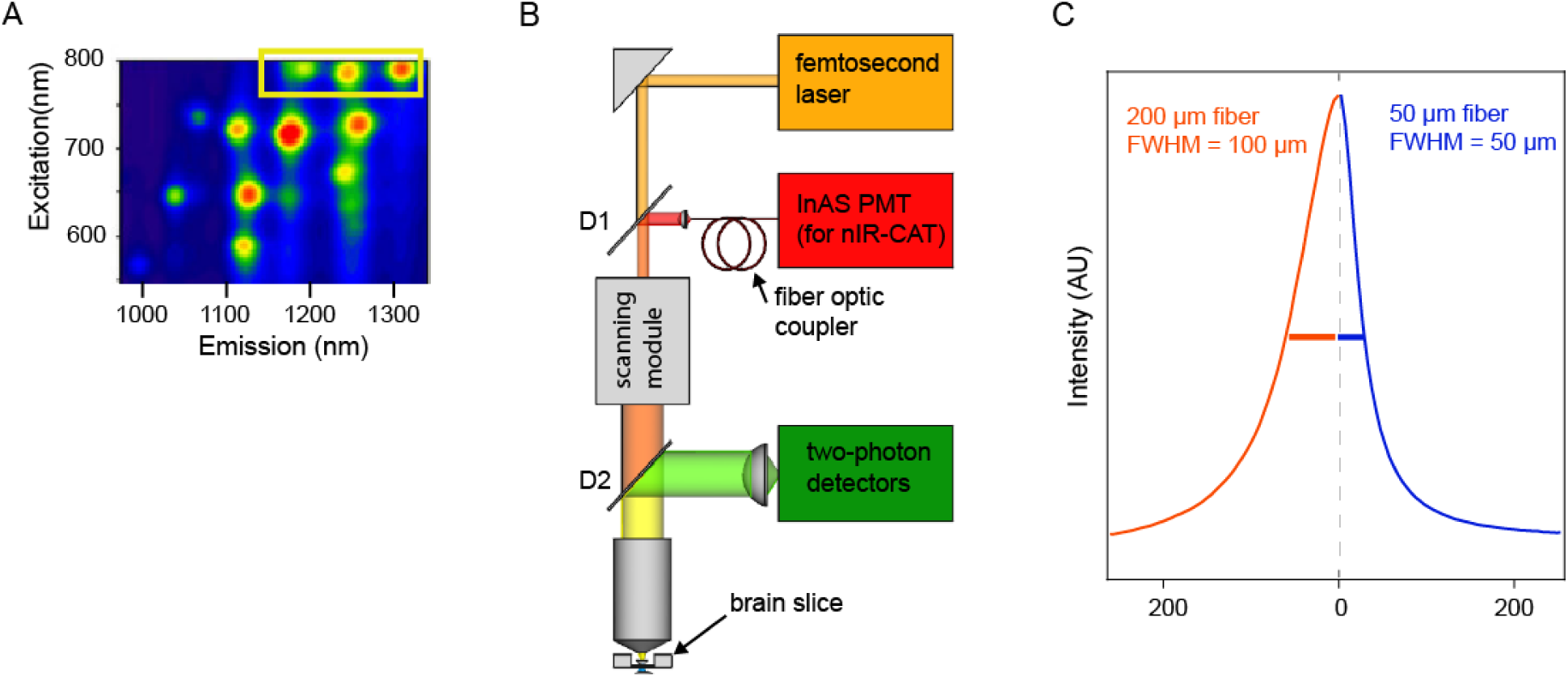
Integrating an infrared detection pathway for imaging of nIRCat fluorescence. **A.** Fluorescence map for nIRcat. Yellow rectangle marks three carbon nanotube chiralities whose excitation overlaps with the 2-photon excitation of GFP. **B.** Microscope schematic. A one-photon infrared (IR) detection arm (red) was added to a scanning two photon microscope. Dichroics: D1: custom dichroic is 680-1040 short pass, reflecting > 1100 nm. D2: 680 nm long pass. There is 1300 nm long pass before InAs PMT. **C.** Limited z-section provided by aperture of fiber optic coupler. Fluorescence profiles of IR fluorescence from lead sulfide quantum dot for thicker fiber (red, 200 µm) and thinner fiber (blue, 50 µm). FWHM was 100 µm for thick fiber and 50 µm for thin fiber.

We integrated an indium arsenide (InAs) photomultiplier tube (PMT, Hamatsu H12397A-75), well established for low-noise infrared detection, into a scanning 2P microscope. The InAs nIR PMT was integrated in the descanned path of the microscope, allowing for the simultaneous imaging of the 2P excitation of visible wavelength sensors and the single-photon confocal imaging of the nIRCat sensors (Figure 1B). Specifically, a 1100-nm short-pass dichroic mirror was inserted into the two-photon excitation path immediately upstream of the scanning mirrors. This dichroic transmits the two-photon excitation beam while reflecting the near-infrared (nIRCat) emission. The reflected nIR fluorescence then passes through a 1300-nm long-pass filter to reject residual reflected excitation light before being focused into a single-mode optical fiber. The fiber acts as a confocal pinhole, rejecting out-of-focus fluorescence from above and below the focal plane and guiding the nIR signal to the InAs photomultiplier tube for detection.

To assess the z-sectioning obtained by the optical fiber, we imaged a thin film of lead sulfide quantum dots adhered to a glass slide using both a 50 µm thick optical fiber and a 200 µm thick optical fiber (Figure 1C). We estimated the effective axial resolution from the FWHM of the z-intensity profile of the resulting image stack. For thick fibers, this corresponds to 100 µm while for the 50 µm fibers it was 50 µm. This limited confocality allows for integration of IR signal over the full range of nIRCat sensors in the tissue.

### Dual real-time imaging of nIRCat and GRAB_DA_ in dorsal striatum

To demonstrate the dual imaging capability of our microscope, we simultaneously imaged evoked-dopamine release in acute brain slices using both nIRCats and GRAB_DA_. We prepared coronal slices from mice expressing GRAB_DA3m_ in the dorsal striatum (Figure 2A and Figure 2-1), a region chosen for its dense dopaminergic innervation and lack of norepinephrine inputs (Berridge and Waterhouse 2003) as nIRCats are also sensitive to norepinephrine (Beyene et al. 2019; Del Bonis-O’Donnell et al. 2021). We then incubated slices with nIRCats for 15 minutes before imaging. Using both visible and nIR channels, we observed robust fluorescent signals in both sensors (Figure 2B). We note that the spatial distribution of GRAB_DA3m_ and nIRCat did not completely overlap (Figure 2B and Figure 2-2), likely because GRAB_DA3m_ is expressed on cell membranes (Sun et al. 2018) whereas nIRCats reside in the brain extracellular space.

**Figure 2:**
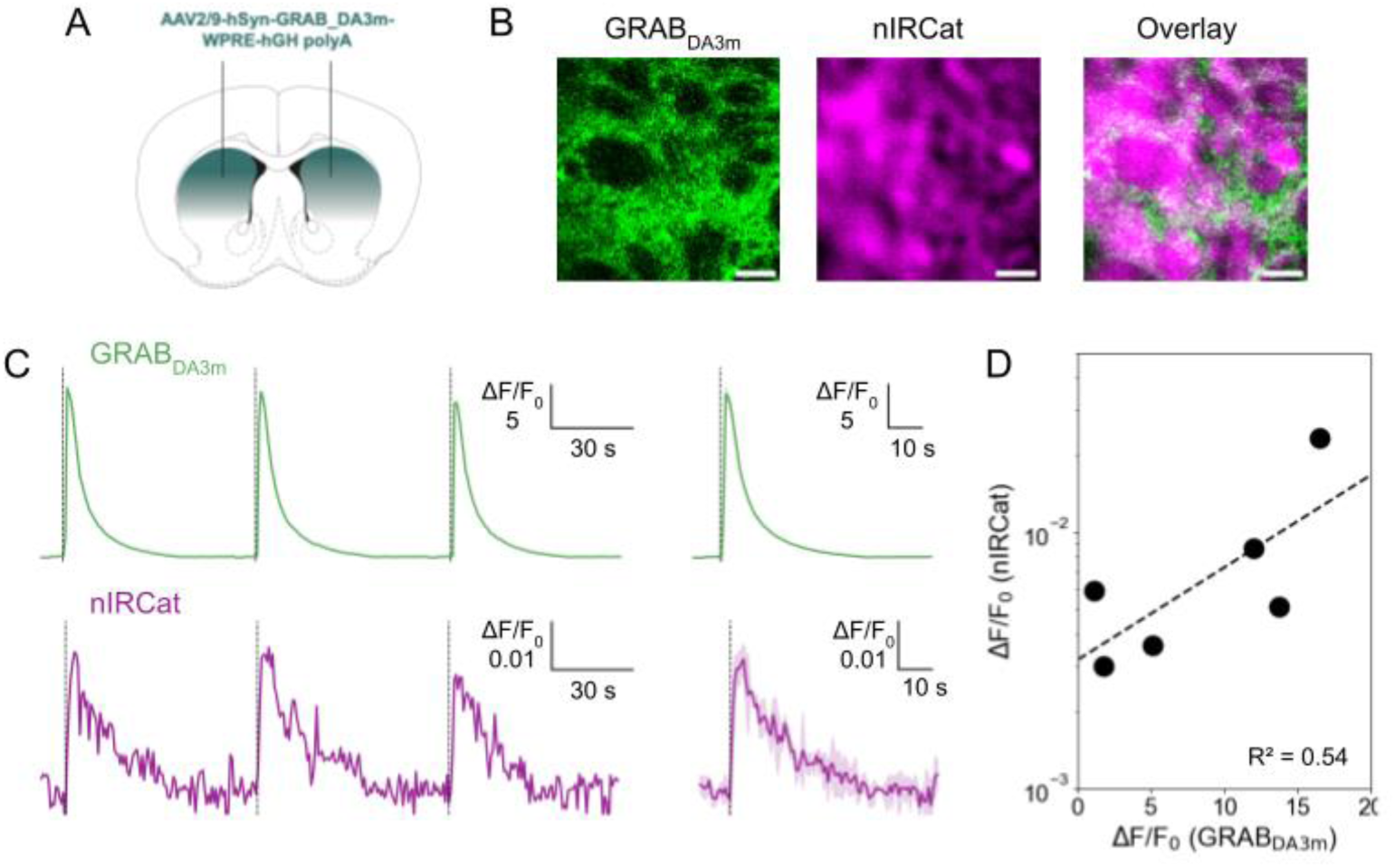
Direct comparison of nIRCat and GRAB_DA3m_. **A.** Scheme of the GRAB_DA3m_ injection sites in the dorsal striatum. **B.** Fluorescence signals from the visible channel (GRAB_DA3m_ ;left), from the nIR channel (nIRCat; middle), and their overlay images. Scale bar denotes 20 µm. **C.** Fractional change (ΔF/F_0_) in fluorescence of GRAB_DA3m_ (top, green) and nIRCat (bottom, purple), integrated from a 74 µm x 74 µm field of view, in response to depolarization via 300 µA electrical stimulation. Individual response (left) and the averaged trace from three repeated stimuli (solid) with SD (shadow) (right). **D.** Comparison of the integrated ΔF/F_0_ between nIRCat and GRAB_DA3m_. The integrated ΔF/F_0_ was obtained from a 74 µm x 74 µm field of view. Each point is the average of three experimental replicates. n = 6 brain slices. Data were fitted with linear regression (dashed line; R^2^ =0.54).

We next electrically evoked dopamine release to assess the microscope’s dual, real-time imaging performance. Depolarization with bipolar stimulating electrodes evoked increases in fluorescence (ΔF/F_0_) in sensors; Both GRAB_DA3m_ and nIRCAT responded reliably to the repeated stimulation (Figure 2C), confirming their reversibility.

Furthermore, the dual imaging capability allowed for a direct comparison of sensor performance: the ΔF/F_0_ of nIRCats was approximately three orders of magnitude smaller than that of GRAB_DA3m_ (Figure 2D), consistent with the sensors’ different affinities, which are discussed in the next section. Note that the ΔF/F_0_ magnitude of nIRCats imaged here was one order smaller than prior reports (Beyene et al. 2019; Del Bonis-O’Donnell et al. 2021), likely because of differences in excitation and detection settings and differences in detector quantum efficiency above 1300 nm. nIRCat signals exhibit a longer decay time (Figure 2C and 2-3), likely reflecting their volumetric sampling of dopamine throughout the extracellular space, combined with the more diffuse optical sectioning in our one-photon imaging. In addition to electrical stimulation, both sensors showed robust responses to short applications of potassium (K^+^) (Figure 2-4). To our knowledge, this study provides the first direct comparison between GRAB_DA3m_ and nIRCat performance under identical experimental conditions.

### nIRCat’s lower affinity enables spatial mapping of high dopamine release sites in dorsal striatum

The dorsal striatum is densely innervated by dopaminergic neurons, resulting in a high extracellular dopamine concentration upon depolarization. Although the high affinity of GRAB_DA3m_ is ideal for many applications, including *in vivo* imaging, it presents a limitation in dopamine-rich regions where the sensor can become saturated, compromising its ability for spatially resolved analyses. Note, GRAB_DA_ sensors with differing affinities have been introduced to address this issue (Zhuo et al. 2024), but their Kd is still in nM range. In contrast, the lower affinity of nIRCats allows for identification of localized areas of high dopamine release, including sub-cellular somatodendritic release as reported earlier (Bulumulla et al. 2022)..

To identify putative dopamine release sites, we applied a 6 x 6 µm grid mask to the images from both GRAB_DA3m_ and nIRCat channels and calculated ΔF/F_0_ for each grid square. Regions of interests (ROIs) were defined as grid squares where peak ΔF/F_0_ exceeded two standard deviations above baseline. Using this approach, nIRCat imaging identified approximately 150 ROIs (out of 361 grid squares) corresponding to “high dopamine spots” (Figure 3A). In contrast, all 361 grids were active in the GRAB_DA3m_ images, consistent with the sensor’s high affinity and therefore saturation (Figure 3C). Next, to assess the sensitivity of these ROIs to changes in dopamine concentration, we applied quinpirole, a D2 dopamine receptor agonist known to decrease dopamine release. Note quinpirole is insensitive to GRAB_DA3m_ because it is based on D1 receptor scaffold and can thus reliably be used to measure dopamine modulation in conjunction with quinpirole (Zhuo et al. 2024). Quinpirole drastically reduced the nIRCat responses, both in overall ΔF/F_0_ (Figure 3A) and in number of identified ROIs (Figure 3B), with a ∼70% reduction across experiments (Figure 3-1). GRAB_DA3m_ exhibited a modest (∼4 %) reduction in overall ΔF/F_0_ (Figure 3C) and no significant change in the number of ROIs (Figure 3D), consistent with sensor saturation. These results highlight that the lower-affinity nIRCat sensor can spatially resolve localized dopamine hotspots, providing a useful tool for mapping high dopamine release microdomains and assessing how these patterns change with genetic manipulation (Black et al. 2025) or environmental condition (Mun et al. 2025).

**Figure 3:**
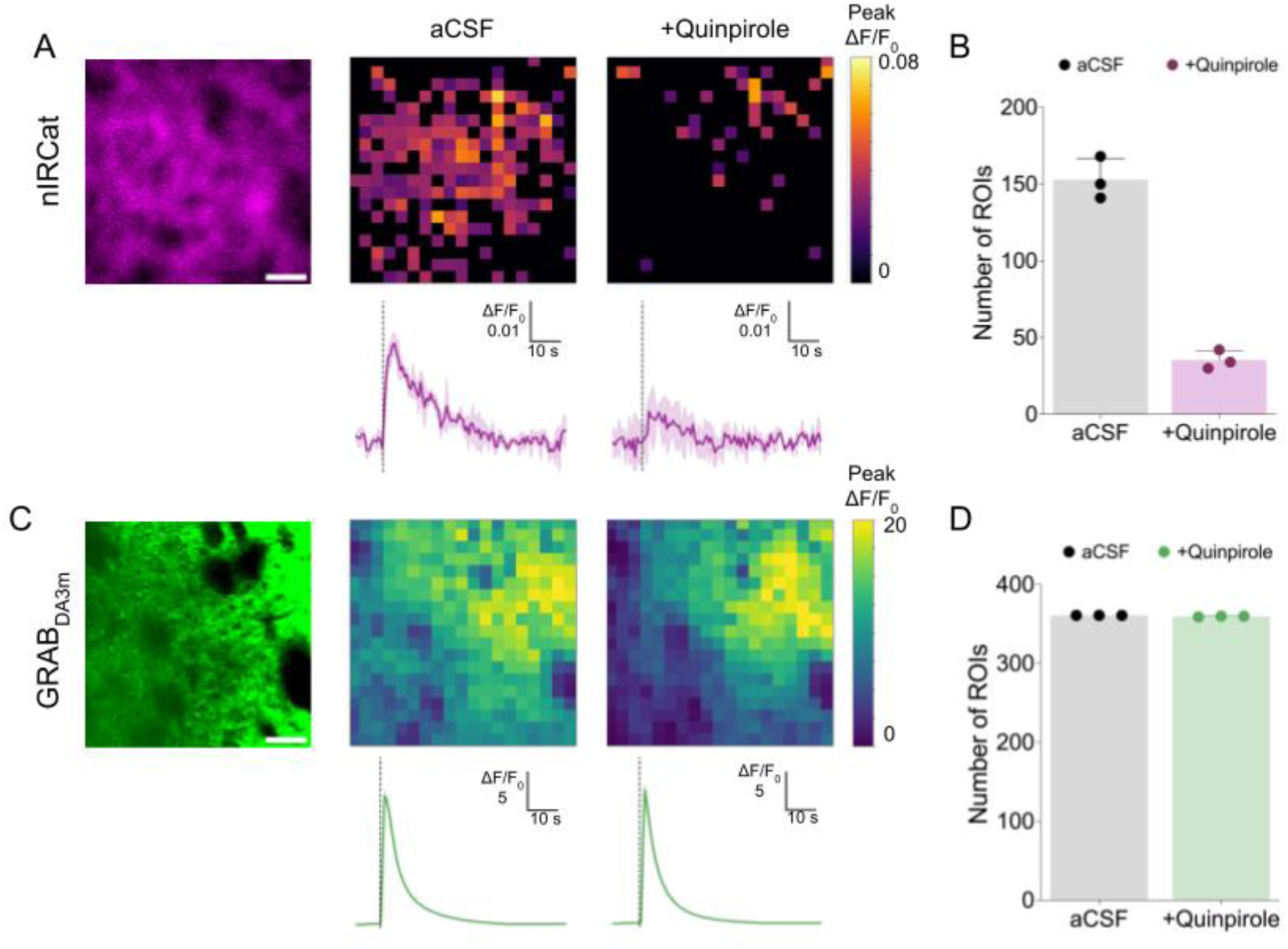
Use of nIRCat to identify localized high dopamine release sites. **A.** (Left) Fluorescent signal from nIRCat. Scale bar denotes 20 µm. 6 µm x 6 µm grid mask was applied to identify regions of interest (ROIs). Heat map of identified ROIs in aCSF (middle) and after 10 µM quinpirole application (right). The color corresponds to peak ΔF/F_0_. (bottom) Integrated ΔF/F_0_ from the entire field of view. Average of n = 3 experiments (solid) with SD (shadow). **B.** The number of identified ROIs in aCSF and after 10 µM quinpirole application. Each data point represents one experiment. n = 3 experiments from 1 brain slice. The error bar is SD. See Figure 3-1 for the summary from n = 3 brain slices. **C.** (Left) Fluorescent signal from GRAB_DA3m_. Scale bar denotes 20 µm. 6 µm x 6 µm grid mask was applied to identify regions of interest (ROIs). Heat map of identified ROIs in aCSF (middle) and after 10 µM quinpirole application (right). The color corresponds to peak ΔF/F_0_. (bottom) Integrated ΔF/F_0_ from the entire field of view. Average of n = 3 experiments (solid) with SD (shadow). **D.** The number of identified ROIs in aCSF and after 10 µM quinpirole application. Each data point represents one experiment. n = 3 experiments from 1 brain slice. The error bar is SD. See Figure 3-1 for the summary from n = 3 brain slices.

**Figure 4:**
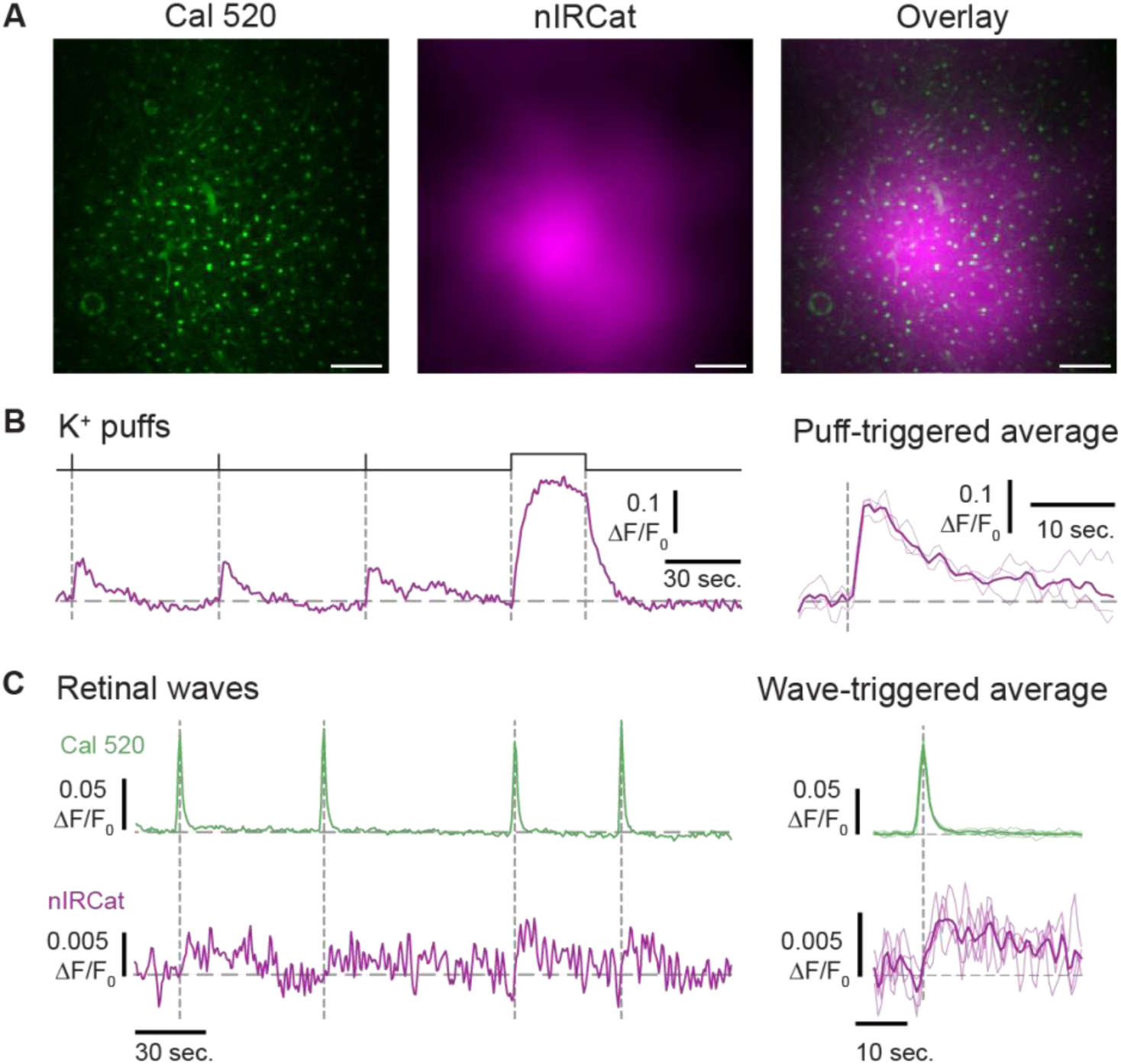
nIRCat response to dopamine release in retina. **A.** Pseudo color images of Cal 520 (green) and nIRCat (purple) in the retina’s inner plexiform layer. Scale bars are 20 µm. **B.** (Left) Fractional change in fluorescence of nIRCat in response to dopamine release evoked by applications of K^+^ (100 mM). Pressure monitor above the traces indicates timing of puffs: three short (1 sec.) and one long (30 sec.) to probe the saturated response. (Right) Puff-triggered average (thick line) and individual trials (thin lines) for the 1 sec. puffs. **C.** (Left) Simultaneous imaging of Cal 520 (green) and nIRCat (purple). Vertical dashed lines mark the peaks of Ca^2+^ transients associated with spontaneous retinal waves. (Right) Wave-triggered averages (thick lines) and signals from individual wave events (thin lines).

### Simultaneous two-photon calcium imaging and nIR dopamine imaging in the developing retina

To demonstrate the applicability of the microscope to other visible fluorophores and other neural circuits, we performed simultaneous two-photon calcium imaging with Cal520 and dopamine imaging with nIRCats in the retina. In the retina, the sole source of dopamine is a sparse population of amacrine cells which play essential roles in both retinal development (Liang et al. 2023; Reis et al. 2007) and adult light adaptation (Roy and Field 2019; Jackson et al. 2012; Goel and Mangel 2021). Although we previously demonstrated that spontaneous retinal waves evoke dopamine release (Arroyo et al. 2016), those experiments were limited by the use of one-photon, FRET-based CNiFER sensors with relatively slow kinetics. As a result, dopamine could only be detected as a bulk signal with limited temporal resolution and no ability to resolve the spatial structure of dopamine release across the retinal tissue. Dopamine release has not been imaged directly from adult retina.

Using our dual-imaging system—which combines near-infrared detection of the dopamine nanosensor nIRCat with two-photon calcium imaging of Cal520—we directly visualized dopamine dynamics in P4–P8 retinas (Figure 4). Brief K⁺ puffs (100 mM, 1 s) evoked robust fractional fluorescence changes in both indicators (Figure 4B), while there was no detectable change with ACSF application or only in carbon nanotubes (Figure 4-1). To assess dopamine release during retinal waves, we repeated these experiments under conditions of spontaneous activity only. Aligning nIRCat signals to spontaneous wave peaks revealed a strong temporal coupling between wave activity and dopamine release (Figure 4C). However, although nIRCat responded robustly to strong depolarization evoked by K⁺ puffs, its lower dopamine affinity resulted in poor signal-to-noise for wave-evoked transients. Hence the lower affinity of the nIRCat limits its ability to detect the lower dopamine concentrations associated with spontaneous retinal waves.

## Discussion

We presented a straightforward approach to introduce a nIR detection capacity to an existing two-photon microscope. This was achieved by introducing a single-mode optical fiber prior to the nIR detector without requiring an additional excitation laser. The fiber-coupled design also provided confocal capability and minimized background nIR signals. This integrated approach allows real-time comparison of dopamine dynamics measured with complementary sensors, as well as simultaneous monitoring of neuronal depolarization and neuromodulator release in intact tissue.

Introducing an nIR channel enabled simultaneous, real-time imaging of visible and nIR fluorescence. Dual-channel imaging is a powerful strategy widely used within the visible spectrum—such as concurrent recording of green and red fluorescence—to reveal complex interactions across cellular and circuit processes (Zhuo et al. 2024). In this study, we extend this capability into the near-infrared range by simultaneously monitoring nIRCat fluorescence alongside dynamically reporting genetically encoded dopamine or calcium indicators in the visible channel. While previous studies have co-imaged nIRCats with neurons labeled with visible fluorephores in culture (Bulumulla et al. 2022; Elizarova et al. 2022), our approach introduces two advances: (1) functional imaging in the visible channel rather than static structural labeling, and (2) confocal capability in the nIR detection by implementing fiber-based coupling, which reduces background and improves optical sectioning for imaging in tissue. Although we focused on dual imaging of nIR and green fluorescence as a proof of concept, this platform is expandable to additional spectral channels. For example, simultaneous recording of acetylcholine activity in green (Jing et al. 2020), calcium dynamics in red (Dalangin et al. 2025), and dopamine release in nIR would allow for multi-transmitter functional imaging within the same preparation.

Multiplexed imaging of sensors with different affinities enables localization of dopamine release sites using a low-affinity sensor, while simultaneously measuring the spatial spread of physiologically relevant dopamine concentrations with high-affinity D1- or D2-based sensors. In addition, GRAB_DA_ is membrane-bound and therefore reports dopamine diffusion near the cell surface, whereas nIRCat is distributed throughout the extracellular volume, allowing measurement of the kinetics of bulk dopamine release. Together, these complementary sensors provide a unified framework for resolving both the spatial and temporal dynamics of dopamine signaling across cellular and tissue scales.

Our demonstration of simultaneous dopamine and calcium imaging in the retina showcased another potential niche for nIRCat applications, where the expression of genetically encoded sensors like GRAB_DA_ is technically challenging. Such contexts include tissues or organs where sensor expression is less characterized (e.g., retina, intestine), early developmental stages before robust expression can occur, or non-traditional model organisms lacking fully annotated genomes (e.g., meadow voles (Mun et al. 2025)).

Live imaging of dopamine release in the retina would address several outstanding questions about how dopamine levels are regulated across development and adulthood (Roy and Field 2019). Here we focused on early postnatal development, when dopamine is released in response to spontaneous retinal waves, which provide patterned activity essential for circuit refinement (Arroyo and Feller 2016; Kirkby et al. 2013). Dopamine release in the retina is confined to a sparse population of amacrine cells, though recent findings indicate that retinal ganglion cells (RGCs) might transiently release dopamine in a manner that influences vascular development (Liang et al. 2023). Important open questions remain regarding the relative contributions of circadian influences versus melanopsin-expressing ipRGC-driven activity in regulating dopamine release (Pérez-Fernández et al. 2019; Zhang et al. 2008). Developing approaches for real-time dopamine imaging will therefore be essential for defining the sources, timing, and spatial dynamics of dopamine signaling during these critical stages of retinal maturation.

Although beyond the scope of this work, adapting this microscope for *in vivo* dual-color imaging would be an exciting next step, given the prevalence of two-photon microscopy in *in vivo* applications. While nIRCat use *in vivo* has not yet been reported, recent progress toward integrating them with fiber photometry (Klinger et al. 2025) suggests a promising direction for future development. However, we note that, due to nIRCat’s lower affinity relative to GRAB_DA_, the latter may remain the easier approach for research groups without access to nIR detectors. Nonetheless, the growing interest in imaging within the near- and far-infrared spectral ranges (e.g., nIR-III window (Feng et al. 2021)) underscores the value of pursuing infrared technologies. Towards that end, the approach described here provides a practical means to integrate longer-wavelength detection capabilities without the need to construct an entirely new microscope.

## Acknowledgements

This work was supported by the National Institutes of Health (NIH) EY013528 (M.B.F.), NIH EY019498 (M.B.F.), F.S.C.-H. was supported by (NEI) F31EY028022-03 and K99EY037374. M.B.F. and B.S. were supported by NIH P30EY003176. We acknowledge support of a Burroughs Wellcome Fund Career Award at the Scientific Interface (M.P.L. and N.K.), a Dreyfus Foundation Award (M.P.L.), the Philomathia Foundation (M.P.L.), an NSF CAREER Award 2046159 (M.P.L.), McKnight Foundation Award (M.P.L.), a Simons Foundation Award (M.P.L.), a Moore Foundation Award (M.P.L.), a Heising-Simons Fellowship (M.P.L.), a Brain Foundation Award (M.P.L.), a Polymaths award from Schmidt Sciences, LLC (M.P.L.), Schmidt Science Fellows, in partnership with the Rhodes Trust (N.K.), and EMBO Long-Term Postdoctoral Fellowship (P.M.). M.P.L. is a Chan Zuckerberg Biohub investigator.

