## Supplemental Files for "Adapting a two-photon scanning microscope for simultaneous single-photon imaging of an infrared dopamine sensor"

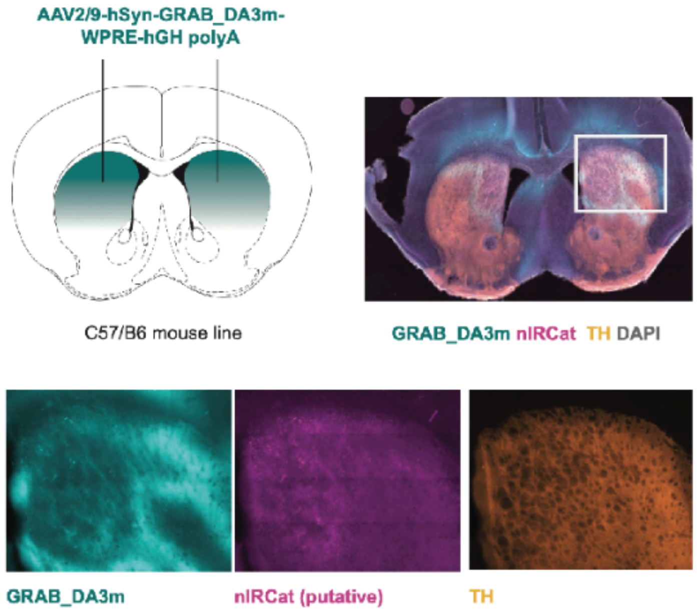


**Figure 2-1. Post hoc histological assessment of $GRAB_{DA3m}$ viral expression in the dorsal striatum.**
A) Schematic of the injection sites
B) Representative striatal slice showing $GRAB_{DA3m}$ viral expression in cyan, putative nIRCat signal magenta (far red wavelength), TH antibody staining in orange.
Bottom line: higher magnification images of the single channels for one of the two hemispheres

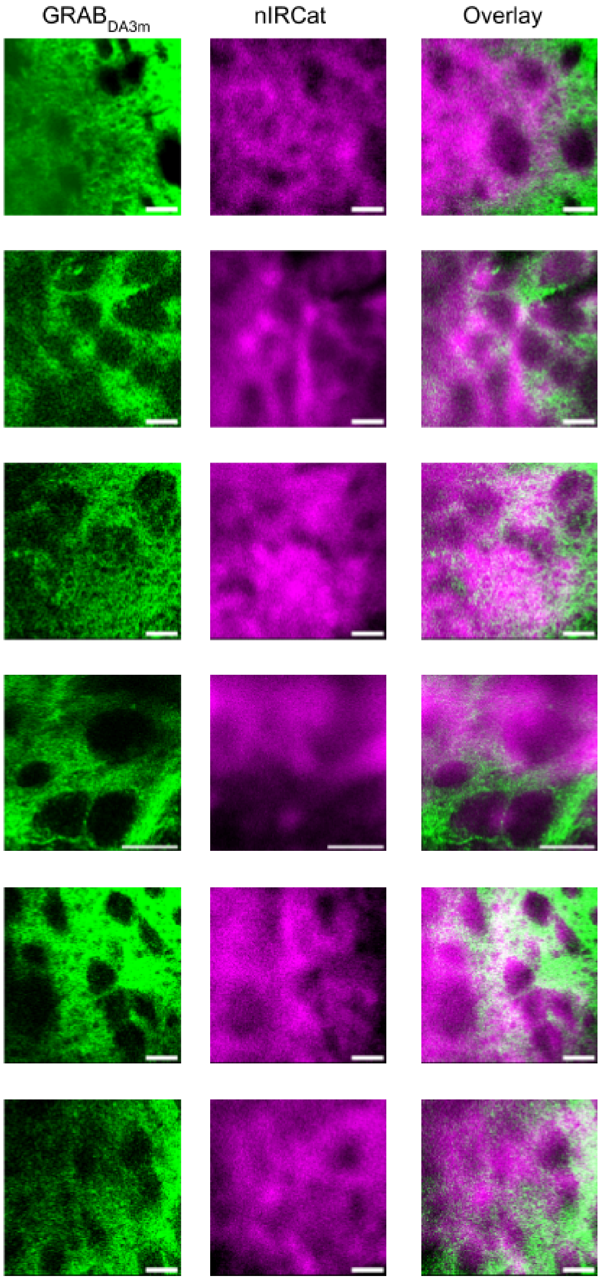


**Figure 2-2. Spatial distribution of GRAB$_{DA3m}$ and nIRCat.** Fluorescent signal from visible channel representing GRAB$_{DA3m}$ (left), from the nIR channel representing nIRCat (middle), and their overlay images from six brain slices. Scale bar denotes 20 μm.

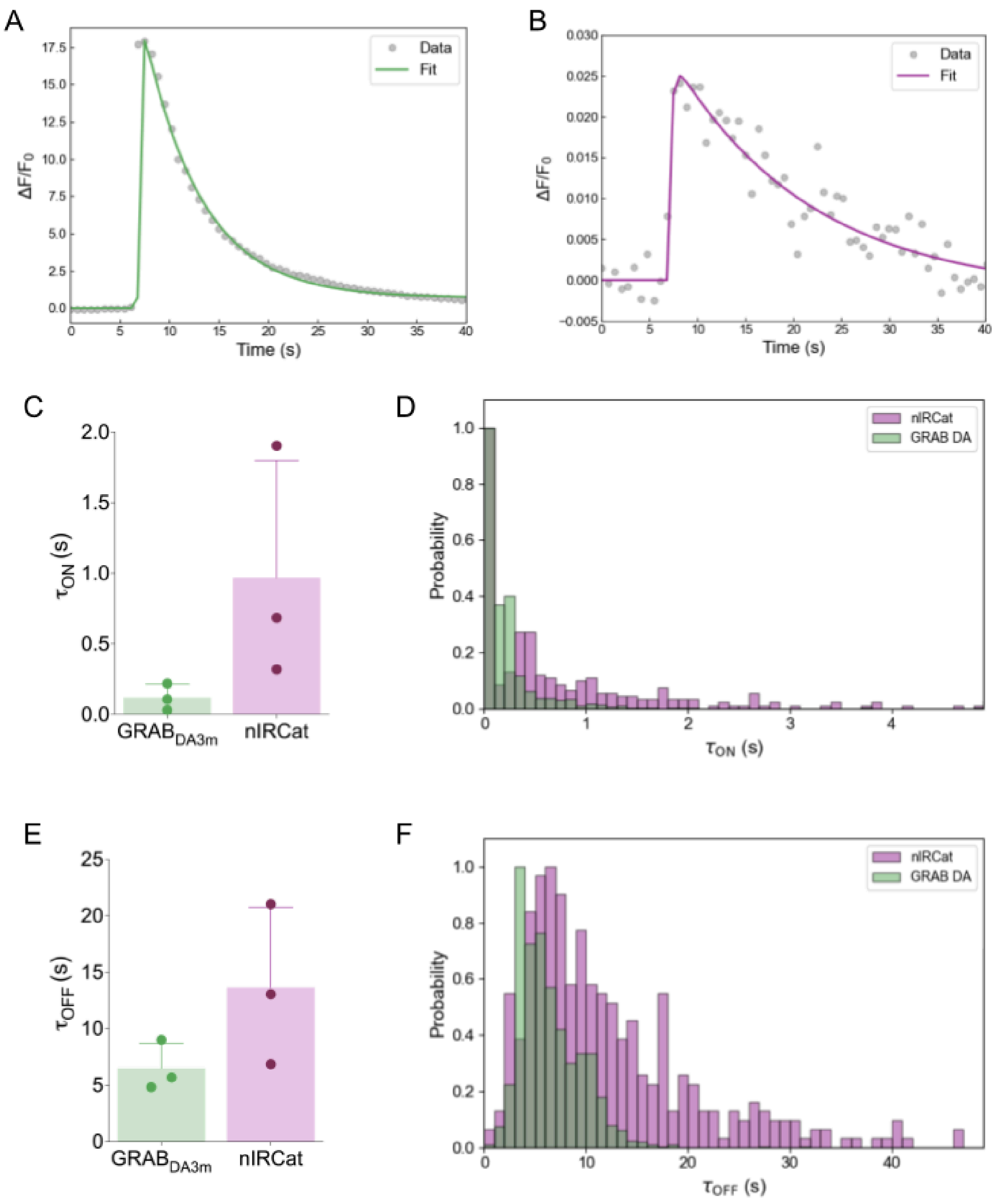


**Figure 2-3. Time constant analysis.**
**A, B**. Example of the fitting procedure for $GRAB_{DA3m}$ (**A**) and nIRCat (**B**) to obtain $\tau_{ON}$ and $\tau_{OFF}$. Obtained $\Delta F/F_0$ data was fitted to

$$\alpha \times \left(1 - exp\left(-\frac{x}{\tau_{ON}}\right)\right) \times \left(exp\left(-\frac{x}{\tau_{OFF}}\right)\right) + \beta,$$

where $\alpha$ is a scale factor, $\beta$ is a constant, $\tau_{ON}$ is the rise time constant, and $\tau_{OFF}$ is the decay time constant. $\alpha$, $\beta$, $\tau_{ON}$, and $\tau_{OFF}$ were fitting parameters.
**C**. Fitted $\tau_{ON}$ for $GRAB_{DA3m}$ and nIRCat from the entire field of view (100 x 100 pixels). Each point is the average of three experimental replicates. n = 3 brain slices. Error bars represent the standard deviation.
**D**. Histogram of $\tau_{ON}$ for $GRAB_{DA3m}$ (green) and nIRCat (pink). 8 x 8 pixel grids were applied to the entire field of view, identified region of interest (ROI) (see Methods for details), and $\Delta F/F_0$ from each ROI was

fitted as shown in A, B. Average from n = 3 brain slices.
**E.** Fitted $\tau_{OFF}$ for $GRAB_{DA3m}$ and nIRCat the entire field of view (100 x 100 pixels). Each point is the average of three experimental replicates. n = 3 brain slices. Error bars represent the standard deviation.
**F**. Histogram of $\tau_{OFF}$ for $GRAB_{DA3m}$ (green) and nIRCat (pink). 8 x 8 pixel grids were applied to the entire field of view, identified region of interest (ROI) (see Methods for details), and $\Delta F/F_0$ from each ROI was fitted as shown in A, B. Average from n = 3 brain slices.

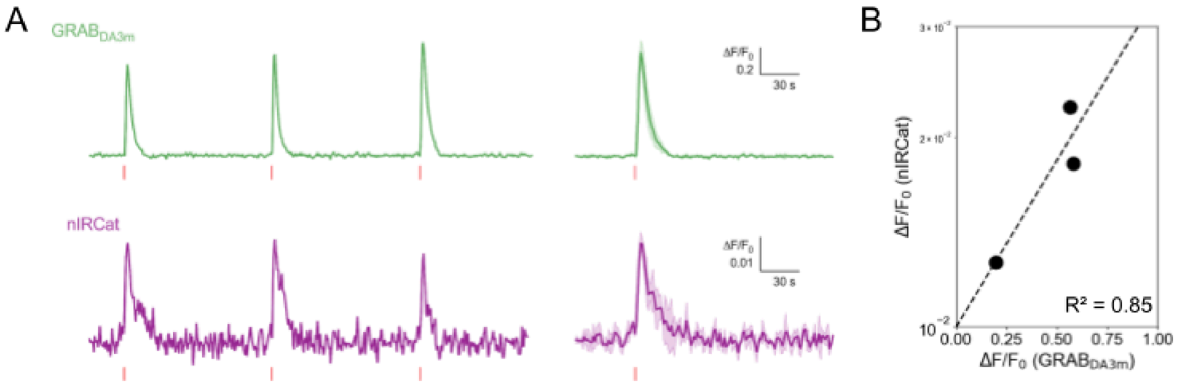


**Figure 2-4. Potassium puff experiments.**
**A.** Fractional change ($\Delta F/F_0$) in fluorescence of $GRAB_{DA3m}$ (top, green) and nIRCat (bottom, purple) recorded in the coronal slice of the dorsomedial striatum in response to depolarization via potassium application (100 mM, 1 s). Individual response (left) and the averaged trace from three repeated stimulation (right).
**B.** Summary of $\Delta F/F_0$ of nIRCat to $GRAB_{DA3m}$. Each point is the average of three experimental replicates. n = 3 brain slices. Data was fitted with linear regression (dash line), and $R^2$ value was 0.85.

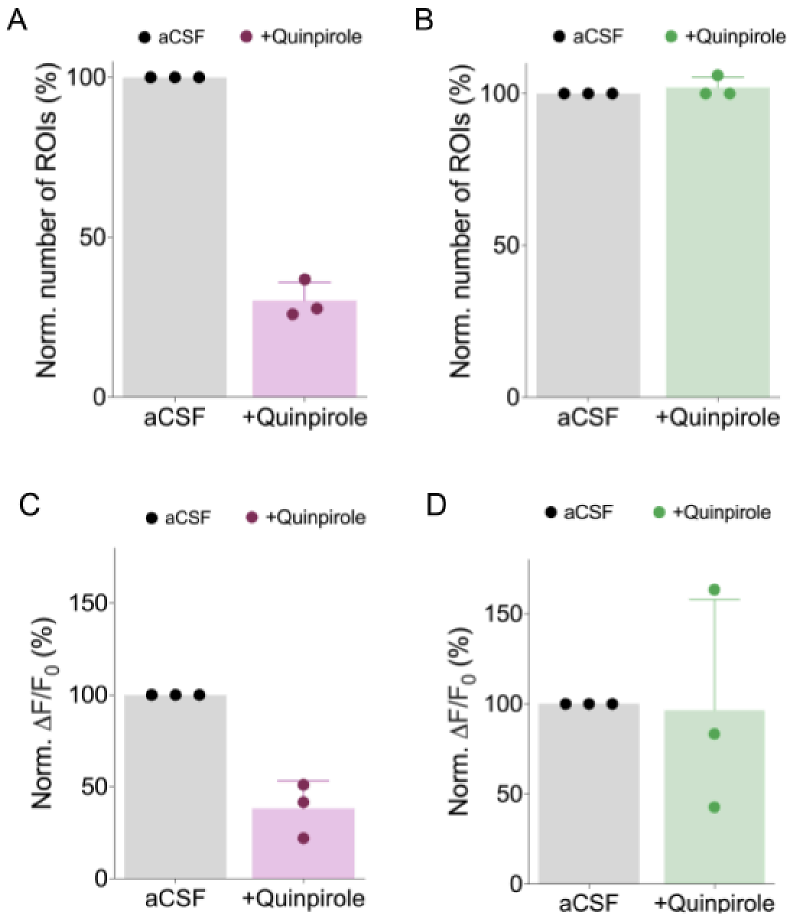


**Figure 3-1. Quinpirole experiments summary.**
**A-B.** The number of ROIs in nIRCat channel (**A**) and $GRAB_{DA3m}$ (**B**) normalized to that of before quinpirole application from 74 µm x 74 µm field of view. Each data point is the average of three experimental replicates. n = 3 brain slices. Error bars represent the standard deviation.
**C-D.** $\Delta F/F_0$ from the entire field of view (74 µm x 74 µm) in nIRCat channel (**C**) and $GRAB_{DA3m}$ (**D**) normalized to that of before quinpirole application. Each data point is the average of three experimental replicates. n = 3 brain slices. Error bars represent the standard deviation.

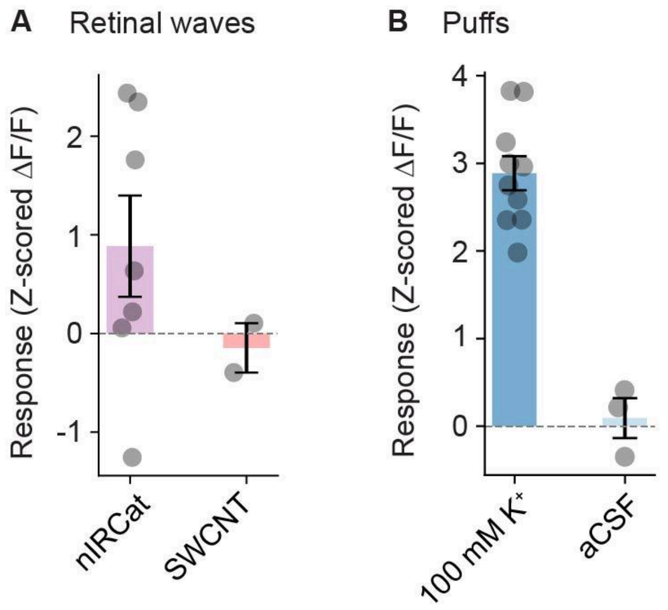


**Figure 4-1. Retinal experiments summary.**
**A.** Responses to retinal waves measured after loading with dopamine-sensitive near-infrared catecholamine sensor (nIRCat) or single-walled carbon nanotubes (SWCNT; a negative control). Each point is an experimental replicate (a retina piece).
**B.** As in A, but for puffs of 100 mM $K^+$ or aCSF (a negative control).
